# Diatoms exhibit dynamic chloroplast calcium signals in response to high light and oxidative stress

**DOI:** 10.1101/2023.08.15.553405

**Authors:** Serena Flori, Jack Dickenson, Trupti Gaikwad, Isobel Cole, Nicholas Smirnoff, Katherine Helliwell, Colin Brownlee, Glen Wheeler

**Affiliations:** The Marine Biological Association, The Laboratory, Citadel Hill, Plymouth, PL1 2PB; Biosciences, College of Life and Environmental Sciences, University of Exeter, Exeter, EX4 4QD, UK

**Author notes:** Corresponding author, 01752 426586.

**Keywords:** diatom, chloroplast, calcium, hydrogen peroxide, *Phaeodactylum*, oxidative stress, signalling

## Abstract

Diatoms are a group of silicified algae that play a major role in marine and freshwater ecosystems. Diatom chloroplasts were acquired by secondary endosymbiosis and exhibit important structural and functional differences from the primary plastids of land plants and green algae. Many functions of primary plastids, including photoacclimation and inorganic carbon acquisition, are regulated by calcium-dependent signalling processes. Calcium signalling has also been implicated in the photoprotective responses of diatoms, although the nature of calcium elevations in diatom chloroplasts and their wider role in cell signalling remains unknown. Using genetically encoded calcium indicators, we find that the diatom *Phaeodactylum tricornutum* exhibits dynamic chloroplast calcium elevations that are distinct from those found in land plants. Chloroplast calcium ([Ca^2+^]_chl_) acts independently from the cytosol and is not elevated by stimuli that induce large cytosolic calcium ([Ca^2+^]_cyt_) elevations. In contrast, high light and exogenous hydrogen peroxide (H_2_O_2_) induce large, sustained calcium elevations in the chloroplast stroma that are not replicated in the cytosol. Measurements using the fluorescent H_2_O_2_ sensor roGFP2-Orp1 indicate that [Ca^2+^]_chl_ elevations induced by these stimuli correspond to the accumulation of H_2_O_2_ in the chloroplast. [Ca^2+^]_chl_ elevations were also induced by the addition of methyl viologen, which acts to generate superoxide within the chloroplast, and by treatments that disrupt non-photochemical quenching (NPQ). The findings indicate that diatoms generate specific calcium elevations in the chloroplast in response to high light and oxidative stress that likely modulate the activity of calcium-sensitive components in photoprotection and other regulatory pathways.

## Introduction

Diatoms are unicellular stramenopile algae that are characterised by their production of a silica cell wall (frustule). They represent one of the most abundant photosynthetic organisms on our planet, playing a major role in both freshwater and marine ecosystems, and are particularly abundant in polar regions and in coastal upwelling systems, which experience substantial changes in light, temperature and nutrient availability. Diatoms therefore require sophisticated cellular signalling mechanisms to sense and respond to their dynamic environment. Recent studies have identified cytosolic calcium signalling mechanisms that mediate the response of diatoms to osmotic, temperature and nutrient stress (Falciatore et al., 2000; Helliwell et al., 2021; Helliwell et al., 2021; Kleiner et al., 2022). However, the role of calcium signalling within other organelles remains unexplored.

In land plants (embryophytes), the chloroplast plays an important role in cellular calcium signalling, either through the modulation of cytosolic calcium elevations or through the presence of Ca^2+^ elevations in the chloroplast itself (Costa et al., 2018). Ca^2+^ concentrations in the thylakoid lumen are 3-5 fold higher than the stroma (Sello et al., 2018). The release of Ca^2+^ from the thylakoid lumen, or its entry across the chloroplast membrane, can result in substantial stromal Ca^2+^ elevations ([Ca^2+^]_chl_, this nomenclature will be used to refer to stromal Ca^2+^). Stromal Ca^2+^ exhibits a strong diel oscillation, increasing during the dark phase (Sai and Johnson, 2002). As many enzymes within the Calvin-Benson-Bassham (CBB) cycle are inhibited by elevated Ca^2+^ (Charles and Halliwell, 1980; Kreimer et al., 1988), it was proposed that the rise in [Ca^2+^]_chl_ plays a direct role in the regulation of photosynthesis. Ca^2+^ is also implicated in the regulation of many other aspects of photosynthesis, suggesting it plays a central role in coordinating signalling pathways within the chloroplast (Hochmal et al., 2016).

Several lines of evidence point to an important role for chloroplast Ca^2+^ signalling in photoprotection. In the green alga *Chlamydomonas reinhardtii,* chloroplast Ca^2+^ signalling modulates photosynthetic electron transport by regulating cyclic electron flow (CEF) around photosystem I, contributing to both the PGRL1/PGR5 and the NAD(P)H dehydrogenase (NDH)-dependent CEF pathways (Hochmal et al., 2015). These responses are mediated by the Ca^2+^ sensor protein CAS, which localises to the thylakoid membrane and forms a complex with Proton Gradient Regulation Like 1 (PGRL1) and Anaerobic Response 1 (ANR1) (Terashima et al., 2012). CAS also regulates the expression of LHCSR3, a light harvesting protein that is required for the dissipation of excess light energy through non-photochemical quenching (NPQ) (Petroutsos et al., 2011). Ca^2+^ therefore regulates the activity of both CEF and NPQ in *C. reinhardtii.* As CAS also regulates the activity of the carbon concentrating mechanism in *C. reinhardtii* (Wang 2016), chloroplast Ca^2+^ signalling plays a central role in coordinating cellular responses to light and inorganic carbon. CAS also plays a role in photoacclimation and stomatal closure in land plants, although its role is less clear. CAS abundance increases in high light in plants and is a target for phosphorylation by the state transition kinase 8 (STN8), which influences CEF (Li et al., 2022). However, CAS is absent from diatom genomes, so the mechanisms of chloroplast Ca^2+^ signalling must differ substantially in this lineage. Several diatom proteins critical for CEF and NPQ possess Ca^2+^-binding domains (Thamatrakoln et al., 2013; Seydoux et al., 2022), suggesting that Ca^2+^ is also important for photoprotection in diatoms but acts via an alternative mechanism.

Chloroplast Ca^2+^ signalling plays an important role in plants and algae, but the dynamics of Ca^2+^ within this organelle remain understudied in comparison with the cytosol. Studies using the bioluminescent calcium reporter aequorin have demonstrated that stimuli that induce [Ca^2+^]_cyt_ elevations, such as oxidative stress and salt (0.3 M NaCl) or the bacterial elicitor flg22 also induce [Ca^2+^]_chl_ elevations, although the kinetics and amplitude of the Ca^2+^ elevations differ between the compartments (Manzoor et al., 2012; Nomura et al., 2012; Sello et al., 2016). Chloroplast specific Ca^2+^ elevations have been observed in *Arabidopsis* seedlings exposed to high temperature (Lenzoni and Knight, 2019). These aequorin-based studies have revealed the presence of [Ca^2+^]_chl_ elevations in response to many different stimuli, although they predominately report the response of a population of cells, which can obscure the complexities of Ca^2+^ dynamics within individual cells.

Direct imaging of Ca^2+^ dynamics in chloroplasts can be achieved by targeted expression of fluorescent genetically encoded calcium indicators (GECI), although their use must be carefully controlled to avoid physiological impacts from the excitation light and fluctuations in stromal pH (Loro et al., 2016; Grenzi et al., 2021). Most fluorescent calcium reporters require excitation with light in the visible range, which has the potential to stimulate light-driven cellular responses, including photosynthesis itself. This problem has been overcome by imaging cells with intermittent light (e.g. every 5 s) to greatly reduce photosynthetic activity, although at the trade-off of limiting temporal resolution (Loro et al., 2016). Light-driven increases in stromal pH, driven by translocation of H^+^ into the thylakoid lumen, also have the potential to affect many fluorescent proteins. Earlier reports suggested stromal pH alkalises strongly in the light (from pH 7 to pH 8), although more recent studies using pH-sensitive fluorescent dyes in isolated chloroplasts from pea or *Arabidopsis* indicate a more modest increase from pH 7.3 in the dark to pH 7.6 in the light (Su and Lai, 2017; Aranda Sicilia et al., 2021). pH sensitivity is an important consideration for single wavelength intensiometric calcium reporters, such as R-GECO1, which have shown much promise for highly sensitive measurements of [Ca^2+^]_cyt_ elevations in plants (Keinath et al., 2015). However, substantial oscillations of cytosolic pH in pollen tubes, guard cells or mesophyll cells had no visible effects on cytosolic R-GECO1 (Li et al., 2021). Measurements of cytosolic Ca^2+^ during pH oscillations in *Arabidopsis* pollen tubes showed that jR-GECO1a gave similar results to the pH-insensitive Ca^2+^ sensor yellow cameleon YC3.6 (Guo et al., 2022). R-GECO1 based Ca^2+^ indicators have previously been used to measure Ca^2+^ dynamics in organelles, such as the mitochondria of animal cells, although they exhibited some sensitivity to large increases in pH induced by 30 mM NH_4_Cl (Hou et al., 2017; Kanemaru et al., 2020). Single wavelength intensiometric Ca^2+^ reporters must therefore be used with caution in organelles that exhibit variable pH (Suzuki et al., 2016). YC3.6 is less sensitive to changes in pH because the individual CFP and YFP components of YC3.6 exhibit a similar degree of pH sensitivity, resulting in little change in the CFP/YFP ratio (Keinath et al., 2015). Direct imaging of stromal-localised YC3.6 has been used to successfully demonstrate [Ca^2+^]_chl_ elevations during light to dark transitions in *Arabidopsis* leaves, and revealed complex additional dynamics such as fast Ca^2+^ spikes in single chloroplasts (Loro et al., 2016).

Chloroplast Ca^2+^ signalling has not yet been explored in diatoms. Photosynthetic stramenopiles acquired their plastids via an endosymbiotic association with a red alga and their plastids demonstrate important organisational differences from those found in the Archaeplastida. Diatom plastids are surrounded by four membranes, with the outer membrane connected to the endoplasmic reticulum (ER) (Falciatore et al., 2020). The thylakoids are not organised into grana, but instead form loose stacks of three vesicles containing the fucoxanthin–chlorophyll-protein light harvesting complexes (Flori et al., 2017). Photoprotection in diatoms utilises a highly efficient fast-responding NPQ component that is dependent on a modified xanthophyll cycle involving the interconversion of diadinoxanthin (Ddx) and diatoxanthin (Dtx) (Lavaud and Kroth, 2006). These structural and functional differences suggest that many signalling processes associated with diatom chloroplasts are likely to be unique.

*Phaeodactylum tricornutum* is a genetically amenable pennate diatom that represents the model species for many aspects of diatom biology, although *P. tricornutum* lacks the characteristic silicified frustule found in most other diatoms (Falciatore et al., 2020). *P. tricornutum* was originally isolated from rock pools on the coast of the UK and has subsequently been identified in a broad range of coastal and brackish locations (De Martino et al., 2007). It is therefore likely to experience substantial variability in physical parameters such as light, temperature and salinity within its natural habitat. Initial studies using aequorin demonstrated the presence [Ca^2+^]_cyt_ elevations in response to hypo-osmotic stress, mechanical stimulation, the addition of iron and the diatom aldehyde decadienal (Falciatore et al., 2000; Vardi et al., 2006). More recently, expression of R-GECO1 in *P. tricornutum* and the centric diatom *Thalassiosira pseudonana* has enabled the visualisation [Ca^2+^]_cyt_ elevations in single diatom cells in response to membrane depolarisation, hypo-osmotic stress, cold temperature and the supply of phosphate (Helliwell et al., 2019; Helliwell et al., 2021; Helliwell et al., 2021; Kleiner et al., 2022). Diatoms therefore exhibit robust cytosolic calcium signalling responses to environmental stimuli, but the involvement of the chloroplast in these responses remains unknown.

In this study, we have examined the nature of chloroplast Ca^2+^ signalling in diatoms. We expressed the intensiometric reporters G-GECO1 and R-GECO1 and the ratiometric reporter G-GECO1-mApple in the chloroplast stroma of *P. tricornutum.* Our results show that [Ca^2+^]_chl_ acts independently of [Ca^2+^]_cyt_ and that both high light and oxidative stress induce large sustained [Ca^2+^]_chl_ elevations in diatoms. By expressing fluorescent reporters for H_2_O_2_ we demonstrate that [Ca^2+^]_chl_ elevations coincide with the accumulation of H_2_O_2_ within the chloroplast.

## Materials and Methods

### Strains and culturing conditions

The wild type *P. tricornutum* strain used in this study was CCAP 1055/1 (Culture Collection of Algae and Protozoa, SAMS, Oban, UK). Cultures were maintained in natural seawater with f/2 nutrients (Guillard, 1975). For imaging experiments, cells were acclimated to an artificial seawater (ASW) medium for minimum 10 days prior to analysis. ASW contained 450 mM NaCl, 30 mM MgCl_2_, 16 mM MgSO_4_, 8 mM KCl, 10 mM CaCl_2_, 2 mM NaHCO_3_, 97 µM H_3_BO_3_, f/2 supplements and 20 mM HEPES (pH 8.0). Cultures were grown at 18 °C with a 16:8 light/dark cycle under an irradiance of 50 µmol m^-2^ s^-1^ and used in mid-exponential phase.

### Generation of *P. tricornutum* strains expressing fluorescent Ca^2+^ and H_2_O_2_ reporters

*P. tricornutum* transformed with the cytosolic R-GECO1 Ca^2+^ biosensor (PtR1) was described previously (Helliwell et al., 2019). All other strains were generated in this study. G-GECO1 and R-GECO1 are intensiometric Ca^2+^ reporters that emit green or red fluorescence respoectively (Zhao et al., 2011). roGFP2-Orp1 is a ratiometric redox sensor that is highly selective for H_2_O_2_ over other forms of ROS (Gutscher et al., 2009). To create plasmids for expression of G-GECO1, R-GECO1 and roGFP2-Orp1 in the cytosol, codon optimised genes were synthesised (Genscript, Netherlands, GenBank Accession OR136874-7) and cloned into the pPha-T1 expression vector via *EcoRI* and *BamHI* restriction sites. Additional constructs for chloroplast-localised biosensors were created by inserting the chloroplast-targeting sequence from *OEE1* into the *EcoRI* site of these plasmids (Gruber et al., 2007; Rosenwasser et al., 2014). The G-GECO1-mApple fusion, with the addition of a GGGSGGGS glycine linker between the two proteins and the *OEE1* chloroplast targeting sequence, was synthesised (Genscript) and cloned into the pPha-T1 expression vector via *EcoRI* and *BamHI* restriction sites. Biolistic transformation of *P. tricornutum* was performed as described in (Helliwell et al., 2019).

### Epifluorescence imaging of biosensors

500 µL of cell culture was added to a 35 mm microscope dish with glass coverslip base (In Vitro Scientific, Sunnyvale, CA, USA) coated with 0.01% poly-L-lysine (Merck Life Science UK, Gillingham, Dorset) to promote cell adhesion to the glass surface. Cells were allowed to settle for 5-20 minutes at room temperature (21°C). R-GECO1 and G-GECO1 were imaged using a Leica DMi8 inverted microscope (Leica Microsystems, Milton Keynes, UK) with a 63x 1.4NA oil immersion objective. A SpectraX LED light source (Lumencor) was used with a 550/15 nm (centre wavelength/bandwidth) excitation filter and 585/40 nm emission filter for R-GECO1 and a 470/24 nm excitation filter and 525/50 nm emission filter for G-GECO1. Images were captured with a Prime 95B sCMOS camera (Teledyne Photometrics, Birmingham, UK) (4 second intervals, 900 ms exposure) using LasX software v.3.3.0 (Leica). Imaging of G-GECO1-mApple used the Leica DMi8 setup but with a PE-300ultra LED light source (CoolLED). Sequential excitation was applied (470/24 nm, 200 ms and 550/15 nm, 200 ms) every 5 s. For continuous light experiments, the excitation shutter was left open after capturing each frame, resulting in continuous illumination by the 470 nm excitation light, although the camera exposure remained the same (200 ms). Imaging of the H_2_O_2_ biosensor roGFP2-Orp1 was performed using sequential ratiometric excitation at 400 nm (390/22) and 470 (470/24) nm, with a 525/50 nm emission filter. Oxidants and other chemical stimuli were administered to cells on the microscope by perfusion using a gravity-fed microfluidics setup. Stock solutions of H_2_O_2_, NH_4_Cl, DTT and methyl viologen were prepared in deionised water, before being added to the ASW at the appropriate concentration. The intensity of the excitation light was measured using an S170C Microscope Slide Power Meter (Thor Labs).

### Processing of imaging data

Images were processed using LasX software (Leica). The mean fluorescence intensity (F) within a region of interest (ROI) encompassing each cell or chloroplast was measured over time. Background fluorescence was subtracted from all cellular F values. The change in the fluorescence intensity of G-GECO1 and R-GECO1 was then calculated by normalizing each frame to the initial value (F/F_0_). Ca^2+^ elevations were defined as any increase in F/F_0_ above a threshold value (>1.5), with sustained Ca^2+^ elevations defined as events where F/F_0_ was greater than 1.5 for >10 s. For G-GECO1-mApple the fluorescence ratio was determined after background subtraction (F_GG/mA_). For roGFP2-Orp1 the fluorescence ratio was determined following excitation at 400 and 470 nm (F_400_/F_470_). The maximum and minimum oxidation states of roGFP2-Orp1 were determined using 1 mM H_2_O_2_ and 1 mM DTT respectively.

### Fluorescent plate reader assays to measure Ca^2+^ and H_2_O_2_

Analyses were performed using a CLARIOstar Plus fluorescence plate reader (BMG LabTech, Aylesbury, UK). G-GECO-mApple was measured using excitation filters at 470/15 nm and 555/20 nm, with emission filters at 515/20 nm and 610/40 nm. roGFP2-Orp1 was measured using excitation filters at 400/15 nm and 475/15 nm, with emission at 515/15 nm. Background fluorescence (seawater with no cells added) was subtracted from all samples. The maximum and minimum oxidation states of roGFP2-Orp1 were determined using 1 mM H_2_O_2_ and 1 mM DTT respectively.

### Chlorophyll fluorimetry

Measurements of Fv/Fm and non-photochemical quenching (NPQ) were made using an AquaPen-C device (Photon Systems Instruments).

### Statistical analysis

Graphs and statistical analyses were performed using OriginPro (Origin Lab, Northampton, MA). Error bars represent standard deviation. Unless indicated otherwise, imaging experiments were repeated at least three times with independent cultures on different days to ensure reproducibility of the response. Statistical analysis of datasets with more than two groups were performed using an ANOVA followed by a Tukey post-hoc test. Box plots indicate the interquartile range (25-75%) and whiskers show 1.5x interquartile range (IQR). The median (horizontal line) and mean (open square) are also shown.

## Results

### Treatments that induce cytosolic Ca^2+^ elevations in *P. tricornutum* do not induce chloroplast Ca^2+^ elevations

We generated strains expressing calcium indicators targeted to the chloroplast stroma through fusion of G-GECO1 or R-GECO1 to the Oee1 plastid-targeting presequence (Gruber et al., 2007) (Fig 1). In unstimulated control treatments, no cells exhibited [Ca^2+^]_cyt_ elevations, although a small proportion exhibited brief transient [Ca^2+^]_chl_ elevations (3 out of 18 chl-G-GECO1 cells; 2 out of 19 chl-R-GECO1 cells). In land plants, many stimuli that induce [Ca^2+^]_cyt_ elevations, such as oxidative stress and NaCl, also induce rapid [Ca^2+^]_chl_ elevations (Sello et al., 2018). *P. tricornutum* exhibits robust [Ca^2+^]_cyt_ elevations in response to hypo-osmotic shock and the resupply of phosphate (Helliwell et al., 2021; Helliwell et al., 2021). Application of hypo-osmotic shock (75% artificial seawater, ASW) resulted in substantial [Ca^2+^]_cyt_ elevations in most cells (15 out of 20 cells), but did not lead to [Ca^2+^]_chl_ elevations. Similarly, addition of 36 µM PO ^3-^ to phosphate-starved cells caused a transient [Ca^2+^] elevation in 46.7% of cells, but did not cause [Ca^2+^]_chl_ elevations. These findings indicate that [Ca^2+^]_chl_ is not directly influenced by large [Ca^2+^]_cyt_ transients, suggesting that [Ca^2+^]_chl_ can act independently from [Ca^2+^]_cyt_.

**Figure 1:**
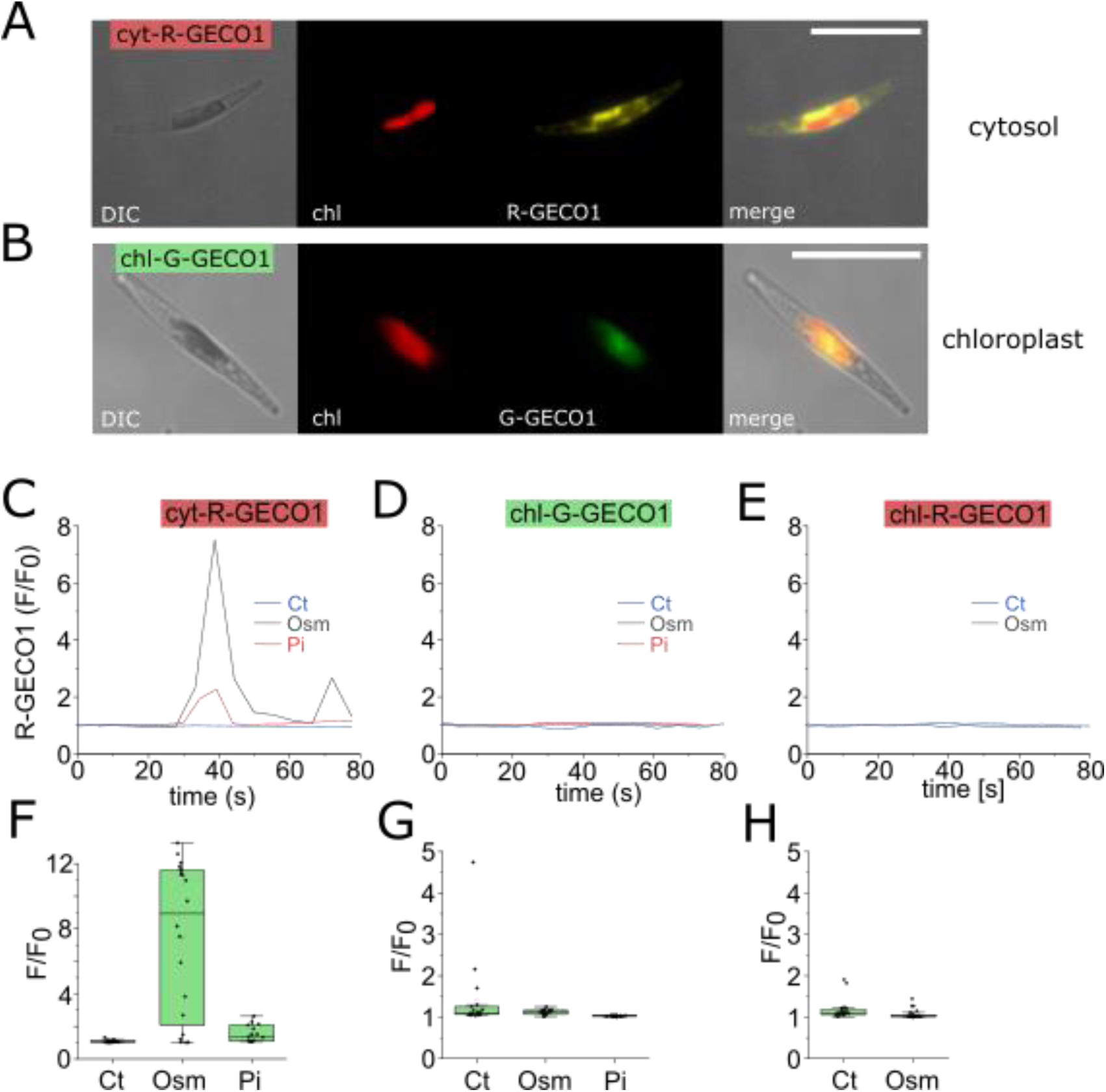
Cytosolic Ca^2+^ elevations occur independently from chloroplast Ca^2+^. **A)** Epifluorescent microscopy images of cells expressing R-GECO1 in the cytosol (cyt-R-GECO1). DIC = differential interference contrast, chl = chlorophyll autofluorescence. Bar =10 µm. **B)** Cells expressing G-GECO1 in the chloroplast stroma (chl-G-GECO1). Bar =10 µm. **C)** Changes in cytosolic Ca^2+^ (cyt-R-GECO1) following hypo-osmotic shock (dilute artificial seawater, 75% ASW) or the addition of inorganic phosphate (36 µM, shaded) to P-limited cells (grown at 1.8 uM Pi for 4 d). A representative trace is shown for each treatment. The shaded area indicates the duration of treatment (27-80 s). Each treatment was repeated at least three times on three independent cell cultures. Ct=control, Osm=hypo-osmotic shock, Pi=inorganic phosphate. **D)** Maximal amplitude for cytosolic Ca^2+^ increases, shown as change in fluorescence (F/F_0_). The box plot indicates interquartile range (25-75%), whiskers 1.5 IQR. The median (line) and mean (open square) are also shown. The number of cells is listed in parentheses. **E)** No changes were observed in chloroplast Ca^2+^ (chl-G-GECO1) following hypo-osmotic shock or the addition of inorganic phosphate. **F)** Maximal amplitude for [Ca^2+^]_chl_ increases measured using chl-G-GECO1. Note that spontaneous [Ca^2+^]_chl_ elevations are occasionally observed in untreated control cells. **G)** [Ca^2+^]_chl_ (chl-R-GECO1) following hypo-osmotic shock. **H)** Maximal amplitude for [Ca^2+^]_chl_ increases measured using chl-R-GECO1.

### Light stress induces specific [Ca^2+^]_chl_ elevations that are not observed in the cytosol

Previous studies using fluorescent indicators in plants have used intermittent excitation (e.g. 5 s intervals) at minimal intensity to prevent activation of photosynthesis (measured through the formation of the trans-thylakoid pH gradient) (Loro et al., 2016; Su and Lai, 2017). Our control imaging conditions (intermittent excitation every 4 s), revealed no [Ca^2+^]_cyt_ elevations within a 4 minute period, but identified occasional [Ca^2+^]_chl_ elevations in a small proportion of cells (Fig 2, Supporting Fig S1). Imaging under high light (intermittent excitation every 4 s at a higher intensity, mean irradiance 2365 μmol m^-2^ s^-1^) had a substantial impact on [Ca^2+^]_chl_. 80.5% of chl-G-GECO1 and 47.6% of chl-R-GECO1 cells exhibited a large sustained (>10 s) increase in [Ca^2+^]_chl_ within the 4 minute observation period (n=41 and 21 cells respectively). No changes were observed in [Ca^2+^]_cyt_. Examination of a strain expressing Ca^2+^ reporters in both compartments (R-GECO1 in the cytosol and G-GECO1 in the chloroplast) confirmed that the large [Ca^2+^]_chl_ elevations induced by excess excitation light do not influence [Ca^2+^]_cyt_ (Supporting Fig S2). The [Ca^2+^]_chl_ elevations were not inhibited by the removal of external Ca^2+^ (Ca^2+^-free ASW + 200 μM EGTA), which does inhibit [Ca^2+^]_cyt_ responses to osmotic, cold and Pi treatments (Supporting Fig S3). The [Ca^2+^]_chl_ elevations therefore do not require Ca^2+^ entry across the plasma membrane and must utilise internal Ca^2+^ stores, such as release of Ca^2+^ from the thylakoid lumen or Ca^2+^ entry across the chloroplast envelope.

**Figure 2:**
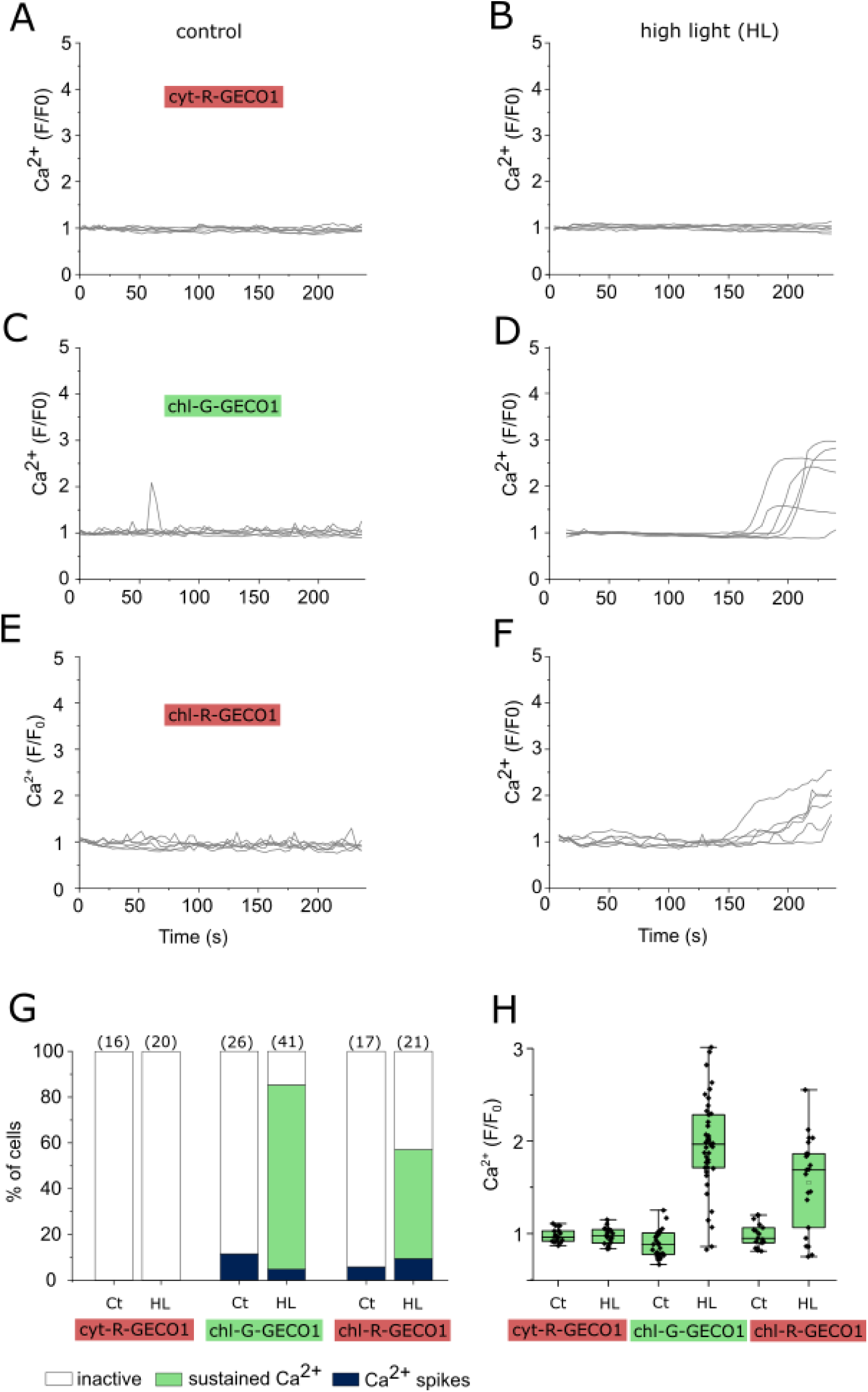
High light causes an increase in chloroplast Ca^2+^. Effect of excitation light on cells expressing cytosolic (cytR-GECO1) or chloroplast (chlG-GECO1 or chlR-GECO1) localised calcium reporters. 6 representative cells are shown for each treatment. **A)** Relative fluorescence (F/F_0_) in cyt-R-GECO1 cells under control imaging conditions (intermittent excitation every 4 s, mean irradiance 451 μmol m^-2^ s^-1^). **B)** Imaging of [Ca^2+^]_cyt_ under high light conditions (intermittent excitation with higher intensity 470 nm LED, mean irradiance 2365 μmol m^-2^ s^-1^). No [Ca^2+^]_cyt_ elevations were observed. **C)** chl-G-GECO1 cells under control imaging conditions. Note the occasional spontaneous [Ca^2+^]_chl_ elevations. **D)** Imaging under high light conditions leads to large sustained [Ca^2+^]_chl_ elevations. **E)** chl-R-GECO1 cells under control imaging conditions. **F)** chl-R-GECO1 cells show a sustained increase in [Ca^2+^]_chl_ after excitation under high light conditions. **G)** Percentage of cells classed as inactive (no Ca^2+^ elevation), showing Ca^2+^ spikes (Ca^2+^ elevations <10 s), or sustained Ca^2+^ elevations (>10 s) at control imaging conditions (Ct) and high light (HL). The number of cells examined is shown in parentheses. **H)** Ca^2+^ elevations after imaging for 4 minutes. The change in fluorescence (F/F_0_) for all three reporters after 4 minutes is shown, number of cells as in (G).

Continuous illumination of chl-G-GECO1 cells at standard intensity (470 nm mean irradiance 1860 μmol m^-2^ s^-1^) also induced a sustained increase [Ca^2+^]_chl_ within 4 minutes (Fig 3). Closer inspection of these traces revealed complex dynamics of [Ca^2+^]_chl_, with multiple rapid Ca^2+^ transients preceding the eventual sustained [Ca^2+^]_chl_ elevation. Moreover, [Ca^2+^]_chl_ elevations exhibited spatial properties, with distinct differences in the timing of [Ca^2+^]_chl_ elevations within different regions of the chloroplast (Supporting Fig S4). We conclude that excess excitation light, either through intermittent illumination at high intensity or continuous illumination at a lower intensity, leads to a sustained elevation of [Ca^2+^]_chl_. Rapid Ca^2+^ spiking preceding a sustained [Ca^2+^]_chl_ elevation was previously observed in individual *Arabidopsis* chloroplasts, although this occurred following a light to dark transition (Loro et al., 2016).

**Figure 3:**
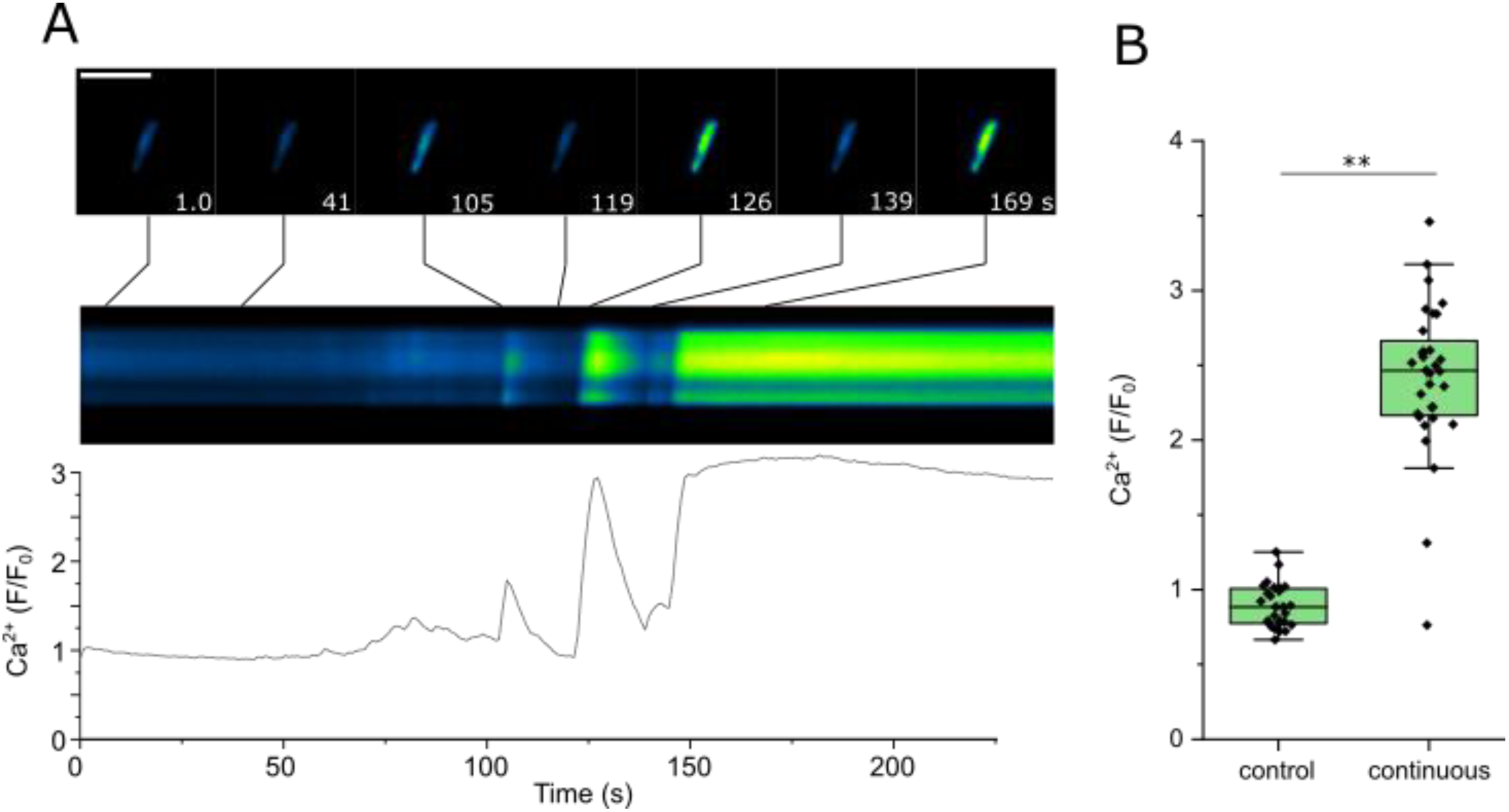
Continuous illumination with blue light results in dynamic [Ca^2+^]_chl_ elevations. **A)** chl-G-GECO1 fluorescence observed under continuous excitation (470 nm at standard intensity, mean irradiance 1860 μmol m^-2^ s^-1^). In the example shown, several transient [Ca^2+^]_chl_ elevations can be observed prior to a sustained [Ca^2+^]_chl_ elevation around 150 s. The trace indicates fold change in fluorescence over time, with the kymograph indicating the change in fluorescence along a longitudinal transect through the chloroplast over this period. Selected individual images from the time course are shown (top panel). Bar = 5 µm. **B) [**Ca^2+^]_chl_ (change in chl-G-GECO1 fluorescence) is shown after 4 minutes for cells imaged under control conditions (intermittent excitation) and continuous illumination. ** = significantly different from control (p <0.01, one-way ANOVA, Tukey post-hoc). n = 27 and 32 cells respectively.

### Ratiometric Ca^2+^ sensors for chloroplast signalling

Single wavelength Ca^2+^ indicators are subject to a series of limitations, such as artefacts caused by cell movement. As the [Ca^2+^]_chl_ elevations in *P. tricornutum* are associated with excess light, changes in stromal pH could also influence R-GECO1 and G-GECO1 fluorescence (emission intensity increases as pH rises). The magnitude of the fluorescence changes in response to [Ca^2+^]_chl_ are likely to be far greater than the pH effect (Li et al., 2021) and we have applied a stringent threshold (F/F_0_>1.5 fold) when categorising [Ca^2+^]_chl_ elevations using single-wavelength Ca^2+^ reporters. However, it is important that all studies of organellar Ca^2+^ dynamics address the issue of reporter pH sensitivity.

The ratiometric Ca^2+^ sensor YC3.6 used previously to monitor [Ca^2+^]_chl_ in plants is largely insensitive to pH because the two fluorophores exhibit a similar pH sensitivity (Loro et al., 2016). Unfortunately, we were unable to successfully target YC3.6 to *P. tricornutum* chloroplasts. We therefore fused G-GECO1 to mApple, which is insensitive to Ca^2+^ but shows some sensitivity to pH (Shen et al., 2014; Rennick et al., 2022), to allow ratiometric imaging of Ca^2+^ with reduced pH sensitivity. We first examined how G-GECO1-mApple responded to changes in pH and Ca^2+^ when expressed in the cytosol. G-GECO exhibited a greater sensitivity than mApple to strong alkalinisation induced by 10 mM NH_4_Cl, although when the fluorophores were used ratiometrically the amplitude of the pH-induced change was significantly reduced (Supporting Fig S5). The maximal increase in cytosolic G-GECO1/mApple fluorescence ratio (F_GG/mA_) induced by strong alkalinisation (10 mM NH_4_Cl) was an order of magnitude lower (59 ± 6 % increase) than typical [Ca^2+^]_cyt_ elevations caused by hypo-osmotic shock (75% ASW) (557 ± 41 % increase). Therefore, although the ratiometric sensor retains a degree of sensitivity to pH, its sensitivity to Ca^2+^ is far greater.

Examination of G-GECO1-mApple in the chloroplasts of unstimulated cells revealed that most cells had a resting F_GG/mA_ between 1.2-1.9 (median value 1.53, n=17 cells) (Fig 4A-B). Imaging chl-G-GECO1-mApple cells for 4 minutes under control conditions (intermittent excitation every 5 s) did not lead to an increase in [Ca^2+^]_chl_ (Supporting Fig S6). We next imaged chl-G-GECO1-mApple cells under control conditions for 60 s (intermittent excitation), before switching to continuous illumination for a further 60 s (470 nm at standard intensity, mean irradiance 4194 μmol m^-2^ s^-1^). mApple exhibited a small increase in fluorescence during the period of continuous light (Fig 4C), which is likely caused by an increase in stromal pH due to activation of photosynthesis by the continuous light (Su and Lai, 2017). G-GECO1 also exhibited a small transient increase in fluorescence at the onset of continuous illumination, followed by a much larger sustained increase that either occurred during the period of continuous light or after the return to intermittent light (n=17) (Fig 4D). These large dynamic changes in G-GECO1 fluorescence were not observed in mApple. Whilst the increase in stromal pH during continuous light treatment could influence G-GECO1 fluorescence, this effect is minor compared to the large sustained changes in fluorescence due to [Ca^2+^]_chl_. Moreover, any pH effect is largely negated when G-GECO1-mApple is used ratiometrically (F_GG/mA_) (Fig 4E). 60 s of continuous illumination caused substantial [Ca^2+^]_chl_ elevations in all cells (n=17). Longer term imaging indicated that [Ca^2+^]_chl_ remained elevated for >10 minutes in all cells before returning to resting values in 62% of cells after 25 minutes (n=21) (Supporting Figure S7).

**Figure 4:**
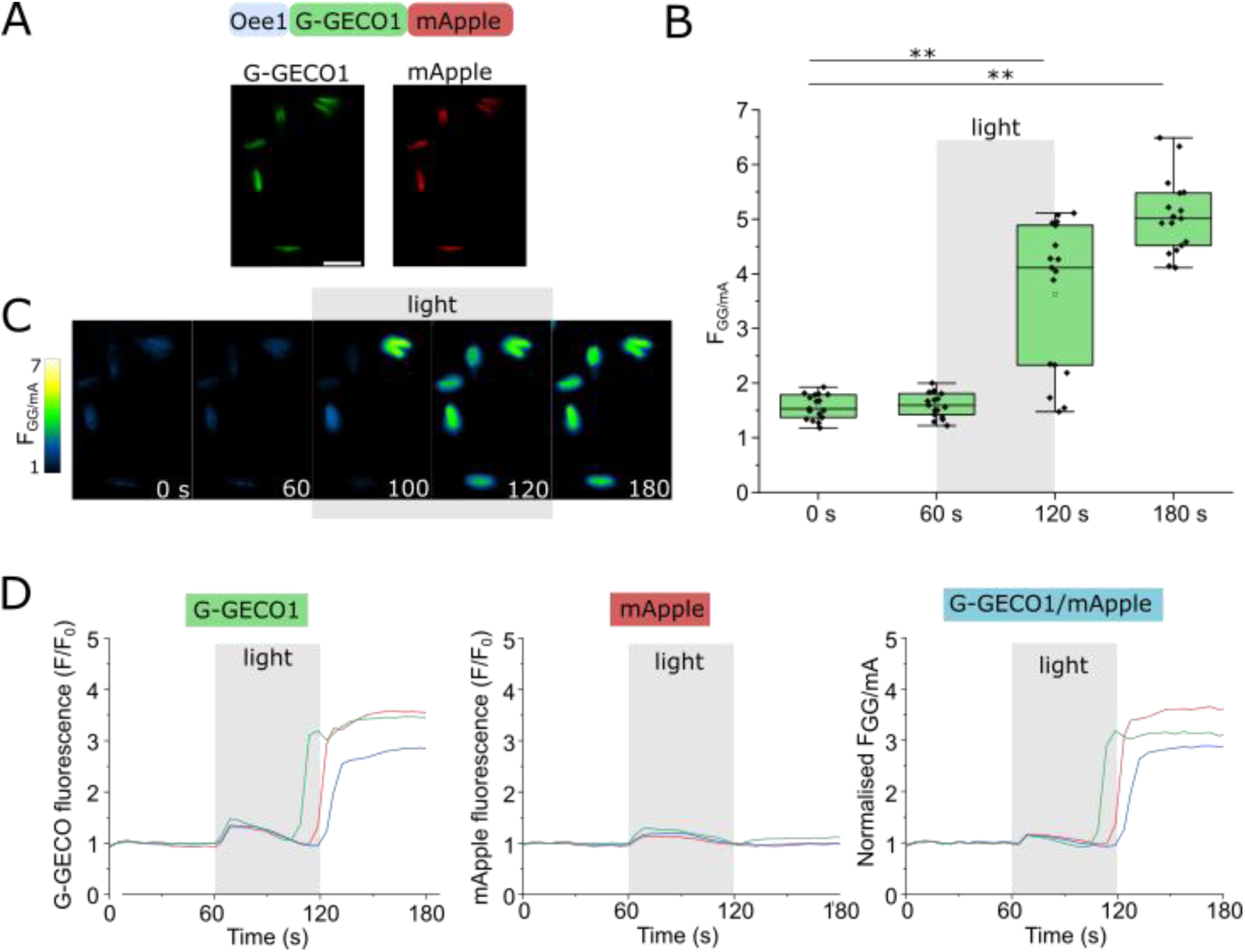
Ratiometric Ca^2+^ indicators demonstrate [Ca^2+^]_chl_ elevations caused by continuous illumination. **A)** Schematic showing the design of a ratiometric Ca^2+^ indicator created by fusing G-GECO1 to mApple. The Oee1 presequence enables targeting to the chloroplast. **B)** [Ca^2+^]_chl_ elevations caused by continuous illumination. Cells were imaged under control conditions (intermittent excitation every 5 s) for 60 s and then exposed to continuous blue light illumination for a further 60 s (mean irradiance 4194 μmol m^-2^ s^-1^). The mean ratio of chl-G-GECO1-mApple fluorescence (F_GG/mA_) increases substantially during the period of continuous illumination and continues to increase even after the return to control imaging conditions. ** = significantly different from initial, p<0.01 (one-way ANOVA, Tukey post-hoc). n=17. **C)** False colour images of chl-G-GECO-mApple cells from the experiment described in (B). Bar=5 µm **D)** Representative traces from individual cells indicating changes in G-GECO1 and mApple fluorescence along with the normalised F_GG/mA_ ratio. mApple is insensitive to Ca^2+^ but shows mild sensitivity to pH. The small increase in mApple fluorescence during the period of continuous light is likely due to an increase in stromal pH. The large sustained increases in G-GECO1 fluorescence are absent in mApple and therefore caused by changes in stromal Ca^2+^.

To further explore the effect of pH on chl-G-GECO1-mApple we perfused cells with 10 mM NH_4_Cl. NH_4_Cl is commonly used as a tool to manipulate cytosolic pH, as NH_3_ readily crosses the plasma membrane and forms NH_4_^+^ in the cytosol, consuming a H^+^ (Boron, 2004). In photosynthetic tissues, NH_4_Cl has an important additional impact, acting to uncouple the photosynthetic electron transport chain from ATP generation, due to alkalisation of the thylakoid lumen and disruption of the trans-thylakoid H^+^ gradient (Dean and Miskiewicz, 2003). Acidification of the thylakoid lumen is important for the activation of non-photochemical quenching (NPQ) in diatoms, and treatment of *P. tricornutum* with 5 mM NH_4_Cl severely inhibits NPQ (Blommaert et al., 2021). Addition of 10 mM NH_4_Cl to chl-G-GECO1-mApple cells resulted in a rapid but relatively small increase in mApple fluorescence that resembled the changes in cytosolic pH caused by NH_4_Cl treatment (Fig 5). Therefore, addition of NH_4_Cl likely alkalises the chloroplast stroma as well as the cytosol. The F_GG/mA_ ratio showed a small transient increase following treatment with NH_4_Cl, followed by a much larger increase in 75% of cells (15 out of 20 cells) that was similar in amplitude to those induced by continuous light. These large [Ca^2+^]_chl_ elevations could result from a direct influence of stromal alkalinisation on Ca^2+^ transport processes, e.g. by modifying the activity of Ca^2+^/H^+^ exchangers, but may also result from disruption of the transthylakoid H^+^ gradient by NH_4_Cl and the subsequent inhibition of NPQ. Alkalinisation of the cytosol by NH_4_Cl did not cause elevations in [Ca^2+^]_cyt_ (Supporting Figure S5), indicating that this response is specific to the chloroplast.

**Figure 5:**
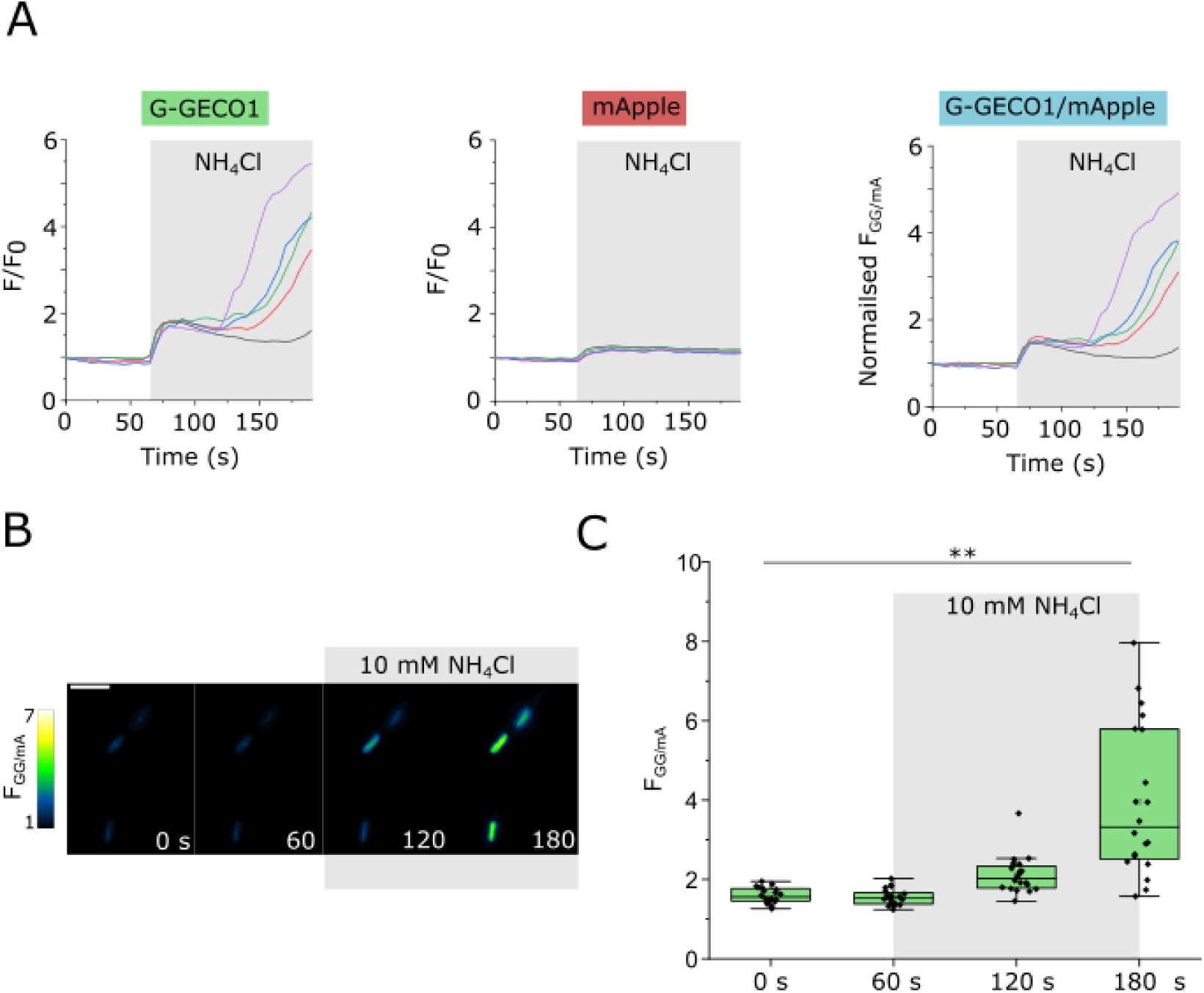
NH_4_Cl induces [Ca^2+^]_chl_ elevations. **A)** Representative traces from individual chl-G-GECO1-mApple cells treated with 10 mM NH_4_Cl. The traces indicate changes in G-GECO1 and mApple fluorescence, along with the normalised F_GG/mA_ ratio. NH_4_Cl causes an increase in stromal pH, which can be observed as a small increase in mApple fluorescence. The large sustained increases in G-GECO1 fluorescence are not seen in mApple. **B)** False colour images demonstrating the large increase in F_GG/mA_ after addition of NH_4_Cl for 2 minutes. Bar=5 µm **C)** Changes in [Ca^2+^]_chl_ (F_GG/mA_) following addition of 10 mM NH_4_Cl. ** = significantly different from initial, p<0.01 (one-way ANOVA, Tukey post-hoc). n=20.

### Chloroplast Ca^2+^ elevations can be induced by oxidants

In photosynthetic organisms, illumination with high light can overwhelm cellular antioxidant defences and lead to an accumulation of reactive oxygen species (ROS) in the chloroplast. NPQ is a major mechanism through which excess excitation energy is dissipated, so inhibition of NPQ can also result in the accumulation of ROS, even at low light intensities (Roach et al., 2015). We hypothesised that the accumulation of ROS may be a common factor in the stimuli that induce [Ca^2+^]_chl_ elevations in *P. tricornutum*. 1 mM hydrogen peroxide (H_2_O_2_) had little impact on [Ca^2+^]_cyt_ (n=9 cells), but triggered substantial [Ca^2+^]_chl_ elevations that were similar in amplitude to those induced by high light or NH_4_Cl (Fig 6A-D). 76.9% of chl-G-GECO1 cells (n=39) and 100% of chl-R-GECO1 cells (n=19) exhibited a sustained [Ca^2+^]_chl_ elevation within a 4 minute period.

**Figure 6:**
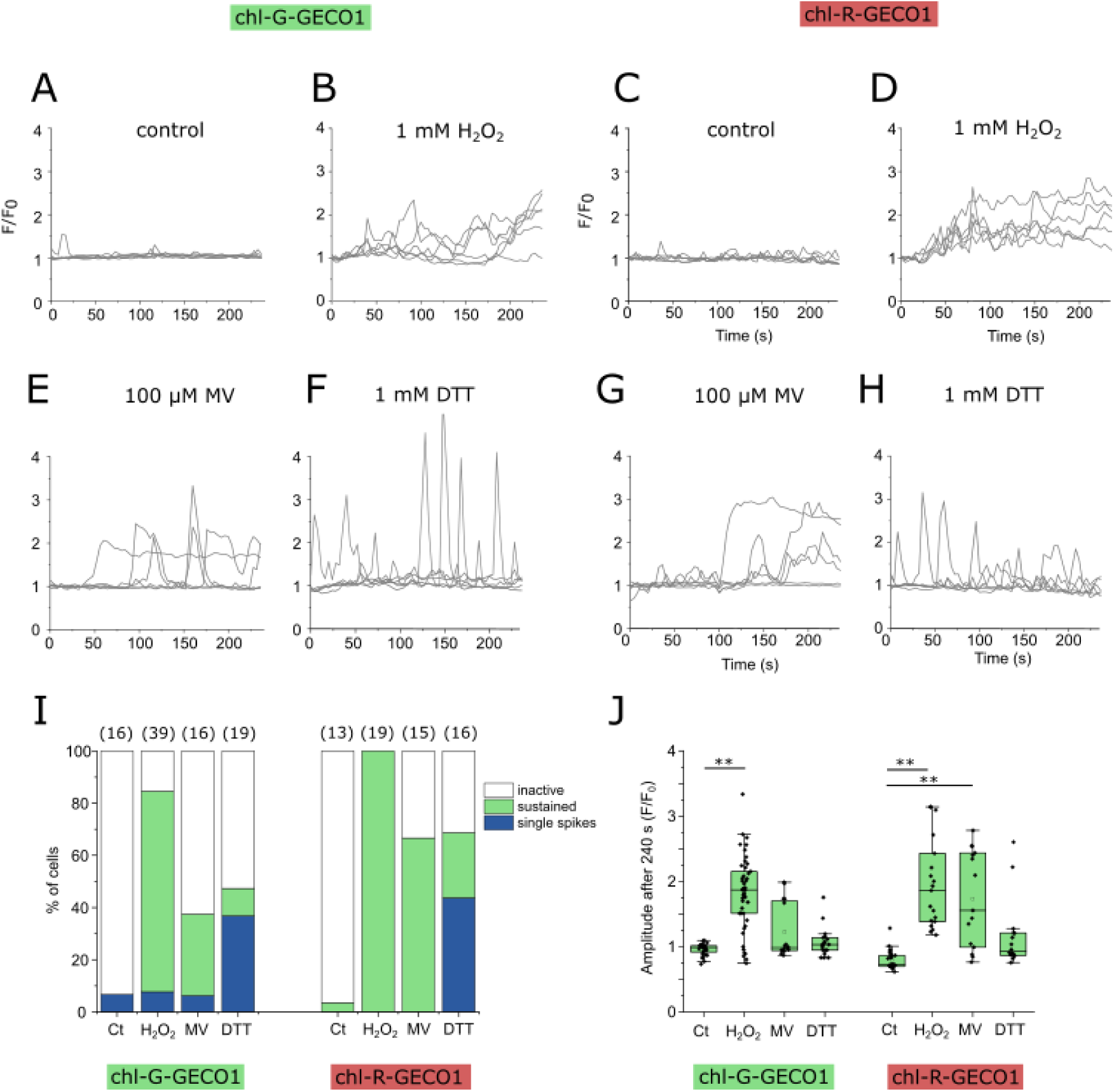
Oxidative stress induces sustained [Ca^2+^]_chl_ elevations. **A)** Control chl-G-GECO1 cells. 6 representative cells are shown for each treatment. **B)** Addition of 1 mM H_2_O_2_ (at time 0 s) leads to a complex [Ca^2+^]_chl_ elevations in all cells. **C)** Addition of 100 μM methyl viologen (MV) also induces [Ca^2+^]_chl_ elevations. **D)** Addition of the reducing agent DTT (1 mM) does not lead to sustained [Ca^2+^]_chl_ elevations, but rapid [Ca^2+^]_chl_ spikes can be observed in many cells. **E-H)** Experiments as described in (A-D) but with chl-R-GECO1 cells. **I)** Percentage of cells classed as inactive (no Ca^2+^ elevation), showing Ca^2+^ spikes (Ca^2+^ elevations that <10 s), or sustained Ca^2+^ elevation (>10 s) for each treatment. The number of cells examined is shown in parentheses. **J)** Change in fluorescence from initial (F/F_0_) for each treatment after 4 minutes. Number of cells as in (I). ** = significantly different from control, p<0.01 (one-way ANOVA, Tukey post-hoc).

Application of lower H_2_O_2_ concentrations (100 μM) to chl-G-GECO-mApple cells resulted in sustained [Ca^2+^]_chl_ elevations in 44.8% of cells within 4 minutes (n=29 cells) (Supporting Fig S8). To measure changes in [Ca^2+^]_chl_ over longer time scales we used a fluorescent plate reader assay to examine chl-G-GECO-mApple cells following treatment with H_2_O_2_. We found that addition of 1 mM H_2_O_2_ led to a significant rise in F_GG/mA_ that could be detected within 10 minutes and remained elevated throughout the 30 minute period (Supporting Fig S9). However, we could not detect a significant increase in [Ca^2+^]_chl_ following addition of 50 μM H_2_O_2_ using this assay.

Methyl viologen (MV) competes with ferredoxin to accept electrons from photosystem I resulting in the formation of superoxide in the chloroplast, which is subsequently converted to H_2_O_2_ by action of superoxide dismutase (Kozuleva et al., 2021). In *Arabidopsis*, MV results in a rapid accumulation of H_2_O_2_ in the chloroplast, with slower and less pronounced elevations in H_2_O_2_ observed in other cellular compartments (Ugalde et al., 2021). We found that 100 μM MV induced substantial [Ca^2+^]_chl_ elevations in *P. tricornutum*, with 31.3% of chl-G-GECO1 and 66.7% of chl-R-GECO1 cells exhibiting a sustained increase in [Ca^2+^]_chl_ (Fig 6E-H). [Ca^2+^]_chl_ elevations can therefore be induced by a treatment that primarily generates ROS within the chloroplast.

We next tested whether addition of a reducing agent (dithiothreitol, DTT), which prevents accumulation of ROS, was able to inhibit [Ca^2+^]_chl_ elevations. Surprisingly, exposure to 1 mM DTT for 4 minutes resulted in a greater frequency of transient [Ca^2+^]_chl_ elevations compared to the untreated control. DTT-treated cells did not exhibit a sustained increase in [Ca^2+^]_chl_, indicating that the nature of the Ca^2+^ signalling response was distinct from those induced by high light, H_2_O_2_ and MV (Fig 6E-H). The results suggest that [Ca^2+^]_chl_ elevations can be induced by perturbation of chloroplast redox state, either to a more oxidised or a more reduced state (Fig 6I-J). In addition to sequestering ROS, addition of DTT will influence many other aspects of redox biology. These include disruption of NPQ through inhibition of the redox-sensitive xanthophyll cycle enzyme violaxanthin de-epoxidase (VDE), which in diatoms catalyses the de-epoxidation of diadinoxanthin (Blommaert et al., 2021). Therefore, two treatments that act via distinct mechanisms to disrupt NPQ (DTT and NH_4_Cl) can both induce [Ca^2+^]_chl_ elevations, albeit with different characteristics.

### Exogenous H_2_O_2_ rapidly increases H_2_O_2_ concentrations within the chloroplast

The sustained [Ca^2+^]_chl_ elevations observed in response to oxidants may be due to direct sensing of the accumulation of ROS in the chloroplast. Using a chloroplast targeted roGFP, which primarily reports the oxidation status of the glutathione pool (*E*_GSH_), van Creveld et al (2015) showed that 80 μM H_2_O_2_ oxidised *P. tricornutum* chloroplasts after 30 minutes. To measure short-term fluctuations in H_2_O_2_ directly, we expressed the ratiometric fluorescent biosensor roGFP2-Orp1 in the cytosol and chloroplast of *P. tricornutum* (Fig 7A). roGFP2-Orp1 is largely insensitive to changes in pH and is highly specific for H_2_O_2_ over other reactive oxygen species (although it can also be oxidised by peroxynitrite) (Nietzel et al., 2019).

**Figure 7:**
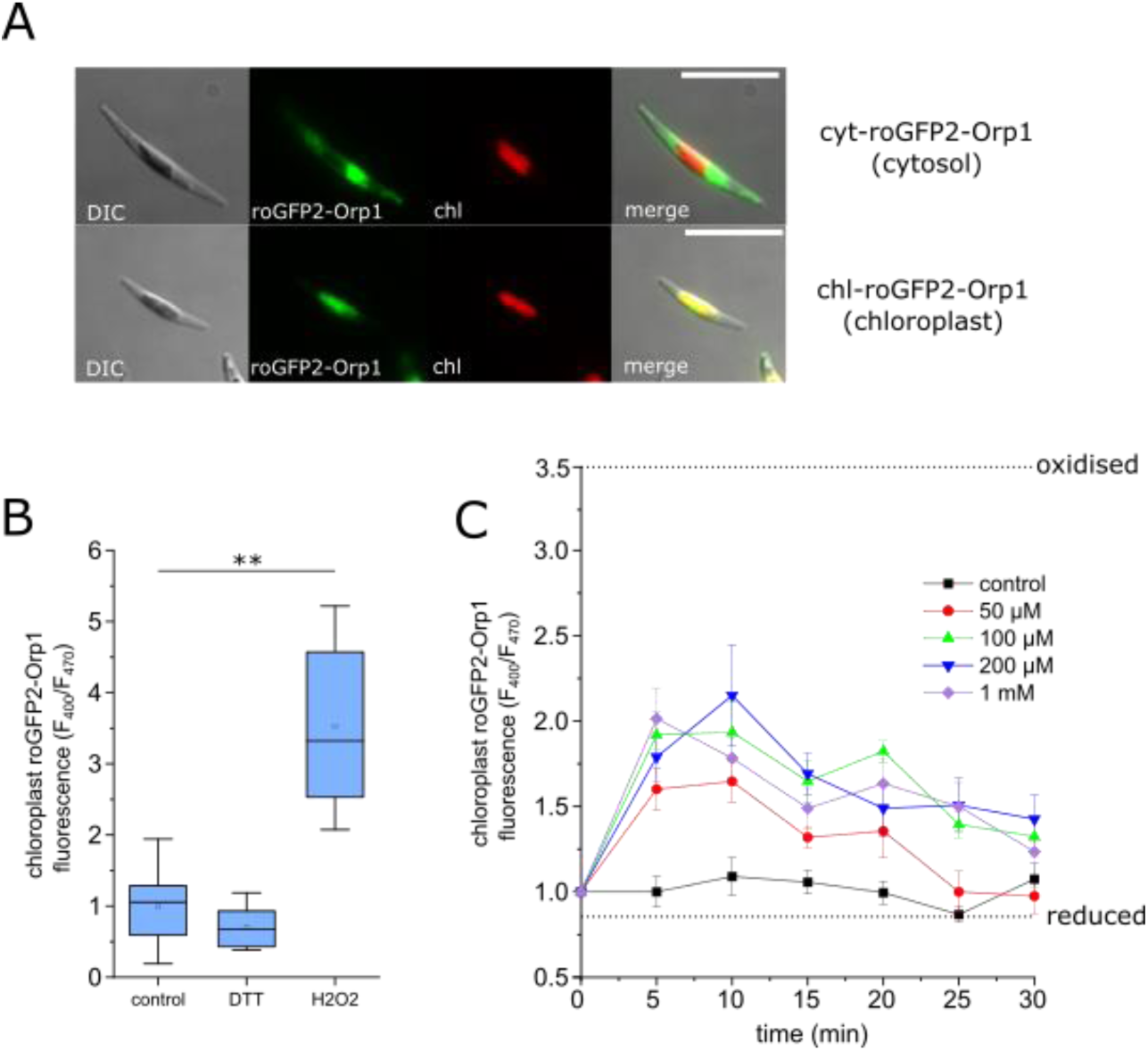
Measurement of chloroplast H_2_O_2_ using roGFP2-Orp1. **A)** Epifluorescent microscopy images of *P. tricornutum* cells expressing roGFP2-Orp1 in the cytosol (upper panel) and chloroplast (lower panel). DIC = differential interference contrast, chl = chlorophyll autofluorescence. Bar = 10 µm. **B)** Determination of roGFP2-Orp1 fluorescence ratio (F_400_/F_470_) when fully oxidised (1 mM H_2_O_2_) or fully reduced (1 mM). Measurements were taken 5 minutes after treatment. ** = significantly different from control, p<0.01 (one-way ANOVA, Tukey post-hoc). **C)** Time course showing the increase in chloroplast H_2_O_2_ in response to the addition of different concentrations of exogenous H_2_O_2_. n=8, error bars represent SE. Dotted lines indicate mean F_400_/F_470_ for fully oxidised and fully reduced probes.

Treatment of *P. tricornutum* cells with 50 μM to 1 mM H_2_O_2_ led to significant oxidation of chloroplast-localised roGFP2-Orp1 within 5 minutes (p<0.05, one-way ANOVA, Tukey post-hoc) (Fig 7B-C). There was a small decrease in F_400_/F_470_ of chloroplast roGFP2-Orp1 following the addition of 1 mM DTT, although this was not significant (p>0.05, one-way ANOVA, Tukey post-hoc). chl-roGFP2-Orp1 remained oxidised in the 100 μM, 200 μM and 1 mM H_2_O_2_ treatments after 15 minutes, but no treatments were significantly different from the control after 30 minutes. Thus, whilst all treatments cause an increase in chloroplast H_2_O_2_, the magnitude and duration of the increase depend upon the strength of the stimulus applied.

### Exogenous H_2_O_2_ has transient effects on photophysiology, but leads to defects in growth

We next examined how longer-term exposure to H_2_O_2_ influenced photosynthesis and growth in *P. tricornutum*. Mizrachi et al. (2019) demonstrated that concentrations of H_2_O_2_ greater than 80 μM caused long-lasting oxidation of the chloroplast within a sub-population of *P. tricornutum* cells that led to their death within 24 h. Three concentrations of H_2_O_2_ (50, 100 and 150 μM) were added to *P. tricornutum* and photosynthetic parameters were measured over 3 h using pulse-amplitude modulated (PAM) fluorimetry. These experiments were performed in cells expressing cyt-roGFP2-Orp1, so that we could directly determine the duration and amplitude of cellular H_2_O_2_ stress at each concentration. Cytosolic H_2_O_2_ was strongly elevated in all treatments 30 min after the addition of exogenous H_2_O_2_, although it returned to resting values after 3 h, with the intensity and duration of the elevation correlating to the concentration of H_2_O_2_ applied (Fig 8A). The photosynthetic efficiency of photosystem II (Fv/Fm) did not change following the addition of 50 µM H_2_O_2_, although there was a significant, but transient, decline in Fv/Fm after treatment with 100 and 150 µM H_2_O_2_ (Fig 8B). Therefore, both cytosolic H_2_O_2_ and Fv/Fm were restored to resting values after 3 h treatment, suggesting that low concentrations of exogenous H_2_O_2_ may only have transient impacts on photosynthesis. However, non-photochemical quenching (NPQ) was significantly elevated after 3 h at 100 μM H_2_O_2_, indicating that the exposure to oxidative stress had a longer lasting impact on photophysiology, even after cytosolic H_2_O_2_ had returned to resting values (Fig 8C). Moreover, growth was severely impaired at all treatments above 50 µM in a dose-dependent manner, indicating that whilst damage to the photosystem was temporary, the oxidative stress experienced was sufficient to inhibit cell division and/or cause cell death (Supporting Fig 10).

**Figure 8:**
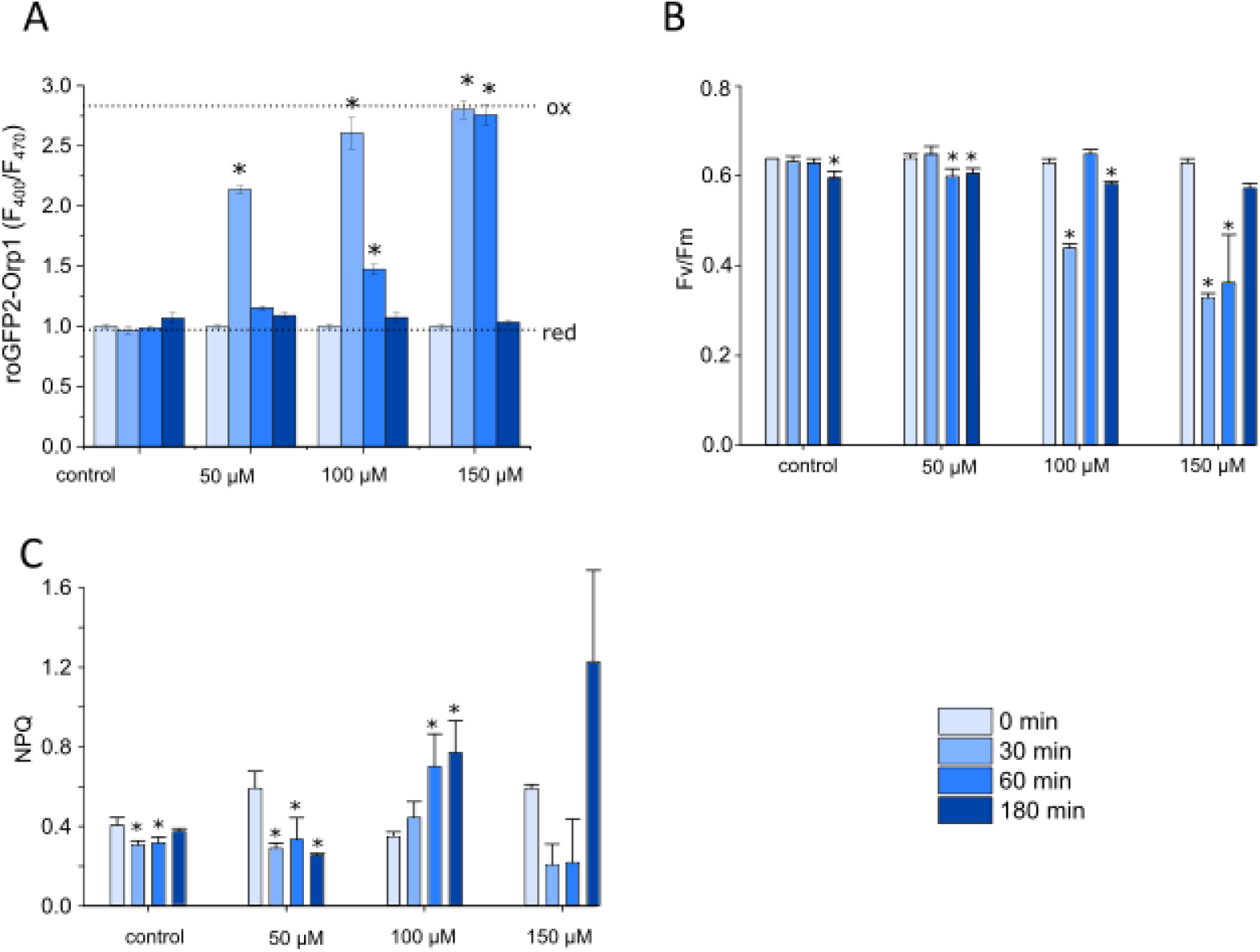
Effect of exogenous H_2_O_2_ on photophysiology. **A)** Cytosolic H_2_O_2_ levels measured using roGFP2-Orp1 after addition of different concentrations of H_2_O_2_ over a 3 h period. n= 3, mean values, error bars represent sd. * = significantly different from initial value for each treatment (P<0.05, one-way ANOVA, Tukey post-hoc). Dotted lines indicate F_400_/F_470_ for fully oxidised and fully reduced probes. **B)** Photosynthetic efficiency of photosystem II (Fv/Fm) in the same cells described in (A). **C)** Measurement of NPQ in the same cells described in (A).

### Light-induced [Ca^2+^]_chl_ elevations coincide with the accumulation of chloroplast H_2_O_2_

Our results suggest that [Ca^2+^]_chl_ elevations induced by high light may be due to an increase in chloroplast ROS. We first examined whether exposure to high light over prolonged periods induced photo-oxidative stress and/or [Ca^2+^]_chl_ elevations in populations of cells within a culture flask. Previous measurements using a chloroplast-localised roGFP indicated that exposure of *P. tricornutum* to 700 μmol m^-2^ s^-1^ for 6 h resulted in only a very slight oxidation of the chloroplast, with much higher irradiances required for oxidation of sub-population of cells (2000 μmol m^-2^ s^-1^ for 6 h) (van Creveld et al., 2015; Mizrachi et al., 2019). In agreement with these findings, we found that exposure of *P. tricornutum* cultures to irradiances of up to 750 μmol m^-2^ s^-1^ for 4 h did not lead to an increase in chloroplast or cytosolic H_2_O_2_ (Fig 9A). We were also unable to detect a significant increase in [Ca^2+^]_chl_ in response to these irradiances using chl-G-GECO1-mApple cells (Fig 9B). The findings indicate that prolonged exposure to irradiances up to 750 μmol m^-2^ s^-1^ does not cause substantial photo-oxidative stress or result in sustained [Ca^2+^]_chl_ elevations in a population of healthy *P. tricornutum* cells.

**Fig 9.**
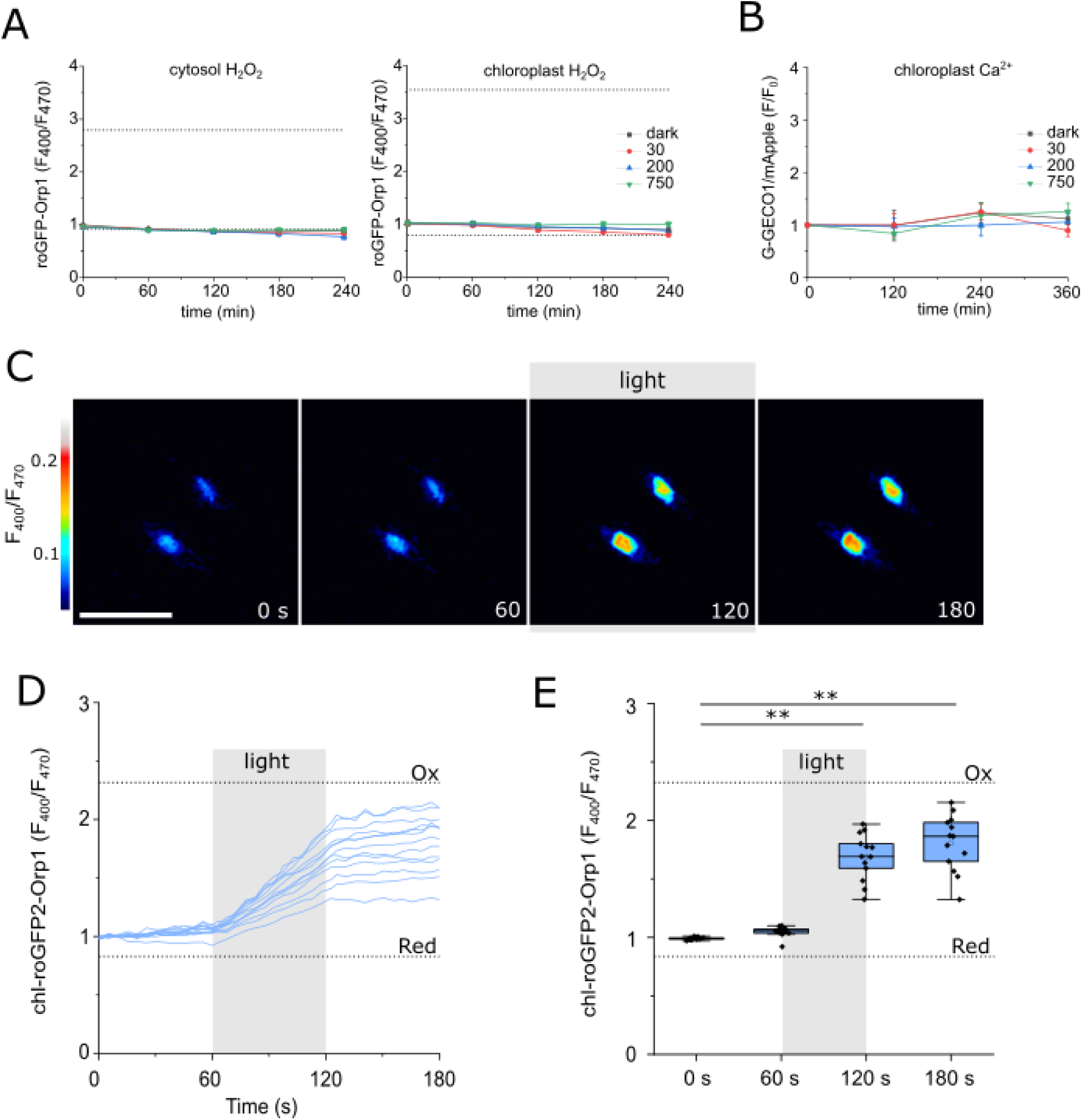
Effect of light on [Ca^2+^]_chl_ and cellular H_2_O_2_. **A)** Measurement of H_2_O_2_ in the cytosol (left panel) and chloroplast (right panel) during exposure to different irradiances over 4 h (dark, 30, 200 and 700 μmol m^-2^ s^-1^). Mean roGFP2-Orp1 fluorescence ratio (F_400_/F_470_) is shown, error bars represent sd, n=5. The dotted lines indicate F_400_/F_470_ for fully oxidised and fully reduced probes. **B)** Changes in [Ca^2+^]_chl_ during exposure of *P. tricornutum* cultures to different irradiances over 6 h. The G-GECO-mApple fluorescence ratio (F_GG/mA_) was measured by a fluorescent plate reader assay and is shown normalised to initial. No significant differences were observed from the 30 μmol m^-2^ s^-1^ control (one-way ANOVA, n=8). **C)** Time course microscopy of individual chl-roGFP2-Orp1 cells exposed to 60 s continuous light at an irradiance that caused [Ca^2+^]_chl_ elevations. Cells were imaged under control conditions for 60 s and then imaged with continuous blue light (470 nm) from 60 s to 120 s. **D)** Representative traces from chl-roGFP2-Orp1 cells showing the accumulation of H_2_O_2_. Dotted lines indicate F400/F470 for fully oxidised (1 mM H_2_O_2_) and fully reduced probes (1 mM DTT). **E)** Box plots showing changes in chloroplast H_2_O_2_ following exposure to continuous light. ** = significantly different from mean initial value, p<0.01 (one-way ANOVA, Tukey post-hoc). n = 12.

We next examined whether brief exposure to the higher irradiances experienced during microscopy led to an accumulation of H_2_O_2_ in the chloroplast of single *P. tricornutum* cells. We imaged chl-roGFP2-Orp1 cells under identical conditions to those used to induce [Ca^2+^]_chl_ elevations in chl-G-GECO1-mApple cells (continuous blue light, 470 nm at 4340 μmol m^-2^ s^-1^ for 60 s) (Fig 4). We found that chloroplast H_2_O_2_ remained stable under standard imaging conditions (intermittent excitation every 5 s) but increased steadily in all cells during continuous illumination (Fig 9C-D). chl-roGFP2-Orp1 remained oxidised following the return to intermittent excitation. The rate at which oxidised roGFP probes return to resting values is determined by the rate at which they are reduced by thiol-based reducing systems in each compartment. In plant cells, the redox potential of the glutathione pool (*E*_GSH_) is the primary determinant of the rate of reduction of oxidised roGFP2-Orp1 (Nietzel et al., 2019). We conclude that irradiances that cause [Ca^2+^]_chl_ elevations also induce a substantial rise in chloroplast H_2_O_2_, supporting the hypothesis that accumulation of H_2_O_2_ plays a direct role in chloroplast Ca^2+^ signalling in diatoms.

## Discussion

Our findings indicate that diatom plastids possess signalling mechanisms that are distinct from those found in the primary plastids of land plants. Multiple stimuli that cause [Ca^2+^]_cyt_ elevations in land plants also cause [Ca^2+^]_chl_ elevations (Sello et al., 2016; Sello et al., 2018). In contrast, we did not observe [Ca^2+^]_chl_ elevations in *P. tricornutum* in response to stimuli that induce [Ca^2+^]_cyt_ elevations. Robust [Ca^2+^]_chl_ elevations in *P. tricornutum* were observed in response to oxidants (H_2_O_2_ and MV), to the reducing agent DTT, to NH_4_Cl and to excess light. Whilst light and H_2_O_2_ also induce [Ca^2+^]_chl_ elevations in land plants, there are important differences. 10 mM H_2_O_2_ induces Ca^2+^ elevations in both the chloroplast and the cytosol of *Arabidopsis* cell cultures (Sello et al., 2016), whereas *P. tricornutum* exhibits Ca^2+^ elevations exclusively in the chloroplast in response to lower concentrations of H_2_O_2_ (100 µM to 1 mM). Light-related [Ca^2+^]_chl_ elevations in land plants occur following a transition from light to dark (Loro et al., 2016). In contrast, [Ca^2+^]_chl_ elevations *P. tricornutum* were caused by exposure to high light intensities. This indicates fundamental differences between the chloroplast Ca^2+^ signalling pathways of land plants and diatoms.

Our evidence suggests that the [Ca^2+^]_chl_ elevations induced by high light in *P. tricornutum* are linked to the formation of ROS in the chloroplast. High light elicited [Ca^2+^]_chl_ elevations at irradiances that generated a rise in H_2_O_2_ in the chloroplast and oxidants were able to cause [Ca^2+^]_chl_ elevations directly. *P. tricornutum* is found in intertidal and coastal locations in temperate regions, where surface irradiance can often exceed 1000 μmol m^-2^ s^-1^ for prolonged periods in summer months (Kolzenburg et al., 2023). As these irradiances can lead to oxidation of the chloroplast redox state in *P. tricornutum* (Mizrachi et al., 2019), diatoms are likely to regularly experience conditions that induce photo-oxidative stress within their natural environment.

The [Ca^2+^]_chl_ elevations induced by NH_4_Cl may also be due to the accumulation of ROS, as this treatment leads to inhibition of NPQ (Blommaert et al., 2021). Disruption of qE (the pH-dependent component of NPQ) in the *npq4* mutant of *Chlamydomonas* leads to a substantial elevation in total cellular H_2_O_2_ compared to wild type cells (Allorent et al., 2013). 1 mM DTT also inhibits NPQ, but this treatment prevents any increase in chloroplast H_2_O_2_. As DTT induced Ca^2+^ spiking rather than sustained [Ca^2+^]_chl_ elevations, transient [Ca^2+^]_chl_ elevations may result from the inhibition of a redox-sensitive process (such as NPQ), whereas sustained [Ca^2+^]_chl_ elevations may result from the accumulation of ROS. These complex signalling events clearly require further dissection, but it seems likely that [Ca^2+^]_chl_ elevations reflect information assimilated from multiple inputs in order to provide a coordinated downstream response.

The link between [Ca^2+^]_chl_ elevations and photo-oxidative stress suggests that chloroplast Ca^2+^ signalling could play a role in the photoprotective response of diatoms. As NPQ is induced in *P. tricornutum* by modest increases in irradiance (e.g. 18 to 135 μmol m^-2^ s^-1^) (Blommaert et al., 2021) that do not cause [Ca^2+^]_chl_ elevations, Ca^2+^ is most likely involved in photoprotective responses to more severe light stress. Several further lines of evidence support a role for chloroplast Ca^2+^ signalling in diatom photoprotection. NPQ is requires establishment of the transthylakoidal proton gradient, which activates the VDE enzyme to promote the formation of diatoxanthin (Dtx) (Lavaud and Kroth, 2006). In plants, the transthylakoidal H^+^ gradient is modulated by the activity of a conserved family of K^+^/H^+^ antiporters, KEA1-3 (Kunz et al., 2014; Wang et al., 2017). A KEA3 homologue also acts as a major regulator of NPQ in *P. tricornutum* (Seydoux et al., 2022). Interestingly, KEA3 from *P. tricornutum* and other diatoms differs from plant KEA3 proteins in that they possess a pair of Ca^2+^-binding EF-hands at the C-terminus. Mutant *P. tricornutum* lines expressing a truncated KEA3 without EF-hands exhibit a similar NPQ phenotype to *kea3* knock-out mutants, indicating that the ability to bind Ca^2+^ is essential for its role in photoprotection (Seydoux et al., 2022). *Arabidopsis* KEA3 is orientated so it mediates H^+^ efflux from the thylakoid lumen (i.e. it lowers the H^+^ gradient when active) with the C-terminal region situated in the stroma (Wang et al., 2017). Assuming PtKEA3 orientates in a similar manner, its EF-hands would be able to detect stromal Ca^2+^ elevations caused by high light and oxidative stress.

Several other Ca^2+^-binding proteins have been identified in diatom chloroplasts (Liu et al., 2022). These include DSP1 (death specific protein), a protein that was first identified as being upregulated during cell death in *Skeletonema costatum* (Chung et al., 2005). DSP proteins localise to the chloroplast and show weak similarity the plant thylakoid-associated proton gradient regulator-5 (PGR5), which contributes to cyclic electron flow (CEF) around photosystem I (Thamatrakoln et al., 2013). All diatom DSP proteins contain a C-terminal pair of EF-hands, suggesting that their activity is also regulated by changes in stromal Ca^2+^ (Chung et al., 2008). DSP proteins appear to play an important role in the responses of diatoms to iron limitation, with *T. pseudonana* lines overexpressing TpDSP1 showing increased growth under iron limitation due to elevated CEF (Thamatrakoln et al., 2013; Hao et al., 2021). The presence of Ca^2+^-binding domains in the key photoprotective proteins KEA3 and DSP1, in combination with the pronounced [Ca^2+^]_chl_ elevations in response to high light, support an important role for chloroplast Ca^2+^ signalling in diatom photoprotection.

Ca^2+^ signalling also plays an important role in programmed cell death (PCD) in diatoms (Vardi et al., 2006). The individual components that mediate PCD are not yet known, although diatoms possess several metacaspases, a family of Ca^2+^-dependent proteases associated with PCD in other eukaryotes. Recent characterisation of a type III metacaspase from *P. tricornutum* (PtMCA-IIIc) revealed that it is co-regulated by Ca^2+^ and redox status (Graff van Creveld et al., 2021). Given that high concentrations of H_2_O_2_ oxidise the chloroplast and lead to cell death in *P. tricornutum* (Mizrachi et al., 2019), it is possible that [Ca^2+^]_chl_ elevations induced by oxidants are also linked to the PCD signalling pathway, particularly as mild H_2_O_2_ treatments result in a similar distribution of chloroplast oxidation and [Ca^2+^]_chl_ elevations in a sub-population of cells. However, the localisation of PtMCA-IIIc has not been reported and PCD caused by the diatom-derived aldehyde decadienal was linked to [Ca^2+^]_cyt_ elevations, rather than chloroplast signalling (Vardi et al., 2006), so it remains to be determined whether [Ca^2+^]_chl_ elevations act in PCD signalling.

Diatom chloroplasts, like those of land plants, can act as autonomous Ca^2+^ signalling organelles. Stromal Ca^2+^ elevations most likely result from release of Ca^2+^ from the thylakoid lumen or the chloroplast ER. Significant recent progress in plants has identified several chloroplast localised Ca^2+^ transporters. *Arabidopsis* BICAT1 and BICAT2 are related to the yeast Ca^2+^ transporter Gdt1 and localise to the thylakoid membrane or the chloroplast envelope respectively (Frank et al., 2019). Disruption of the BICAT proteins leads to substantial changes in the [Ca^2+^]_chl_ transients invoked by light to dark shifts (Frank et al., 2019). In addition, a member of the mitochondrial calcium uniporter family has been shown to localise to the chloroplast in *Arabidopsis* (cMCU), where it contributes to [Ca^2+^]_chl_ dynamics in response to osmotic stress (Teardo et al., 2019). Plastid-localised glutamate receptors GLR3.4 and GLR3.5 also contribute to Ca^2+^ entry from the cytosol (Teardo et al., 2011; Teardo et al., 2015). The entire MCU complex is missing from diatoms (Pittis et al., 2020) and GLRs are also absent from *P. tricornutum* (Verret et al., 2010), so the mechanisms mediating Ca^2+^ entry into diatom plastids are likely to differ substantially from those in land plants.

Our results reveal a highly dynamic Ca^2+^ signalling system within diatom chloroplasts that can act independently of cytosolic Ca^2+^ signalling pathways. [Ca^2+^]_chl_ elevations can be induced by a variety of stimuli linked to photo-oxidative stress, suggesting that chloroplast Ca^2+^ signalling may play an important role in diatom photoprotection. Important distinctions in the nature of chloroplast Ca^2+^ signalling between diatoms and land plants, and in the mechanisms through which Ca^2+^ signals are generated and sensed, suggest that diatoms possess unique signalling mechanisms to regulate photosynthetic function.

## Acknowledgements

The work was supported by an ERC Advanced Grant to CB (ERC-ADG-670390) and a NERC award to GLW (NE/T000848/1). IC was supported by a BBSRC SWBio DTP studentship.

## Author contributions

SF, CB and GLW planned and designed the research. SF, JD, TG, IC and GLW performed experiments and analysed data. SF, NS, KEH, CB and GLW interpreted data and wrote the manuscript.

## Supporting Information

Fig S1 Excitation protocols used during Ca^2+^ imaging.

Fig S2 Simultaneous measurement of Ca^2+^ in the cytosol and chloroplast.

Fig S3 External Ca^2+^ is not required for the high light response

Fig S4 Spatial specificity in [Ca^2+^]_chl_ elevations.

Fig S5 Testing the pH sensitivity of cyt-G-GECO1-mApple.

Fig S6 Imaging chl-G-GECO-mApple under control conditions.

Fig S7 Chloroplast Ca^2+^ recovery after continuous light stress.

Fig S8 [Ca^2+^]_chl_ elevations induced by 100 μM H_2_O_2_.

Fig S9 H_2_O_2_ leads to sustained [Ca^2+^]_chl_ elevations

Fig S10 Longer term impacts of exogenous H_2_O_2_ on growth and photophysiology.

